# EPAC regulates von Willebrand factor secretion from endothelial cells in a PI3K/eNOS-dependent manner during inflammation

**DOI:** 10.1101/2020.09.04.282806

**Authors:** Jie Xiao, Ben Zhang, Zhengchen Su, Yakun Liu, Thomas R. Shelite, Qing Chang, Pingyuan Wang, Alexander Bukreyev, Lynn Soong, Yang Jin, Thomas Ksiazek, Angelo Gaitas, Shannan L. Rossi, Jia Zhou, Michael Laposata, Tais B. Saito, Bin Gong

## Abstract

Coagulopathy is associated with both inflammation and infection, including infection with the novel SARS-CoV-2 (COVID-19). Endothelial cells (ECs) fine tune hemostasis via cAMP-mediated secretion of von Willebrand factor (vWF), which promote the process of clot formation. The <u>e</u>xchange <u>p</u>rotein directly <u>a</u>ctivated by <u>c</u>AMP (EPAC) is a ubiquitously expressed intracellular cAMP receptor that plays a key role in stabilizing ECs and suppressing inflammation. To assess whether EPAC could regulate vWF release during inflammation, we utilized our *EPAC1*-null mouse model and revealed an increased secretion of vWF in endotoxemic mice in the absence of the EPAC1 gene. Pharmacological inhibition of EPAC1 *in vitro* mimicked the *EPAC1*−/− phenotype. EPAC1 regulated TNFα-triggered vWF secretion from human umbilical vein endothelial cells (HUVECs) in a phosphoinositide 3-kinases (PI3K)/endothelial nitric oxide synthase (eNOS)-dependent manner. Furthermore, EPAC1 activation reduced inflammation-triggered vWF release, both *in vivo* and *in vitro*. Our data delineate a novel regulatory role of EPAC1 in vWF secretion and shed light on potential development of new strategies to controlling thrombosis during inflammation.

**Key Point:** PI3K/eNOS pathway-mediated, inflammation-triggered vWF secretion is the target of the pharmacological manipulation of the cAMP-EPAC system.

## Background

Coagulopathy is associated with both severe inflammation and infections, including the coronavirus-induced disease 2019 (COVID-19) ^1-5^. To maintain vascular patency, vascular endothelial cells (ECs) adjust the balance between blood coagulation, bleeding, and fibrinolysis on their luminal surfaces via complicated mechanisms^6-16^. Accumulating evidence has suggested an extensive cross-talk between coagulation and inflammation, whereby inflammation leads to activation of coagulation, and coagulation also considerably affects inflammatory activity^17-19^. Systemic infectious diseases can activate ECs and propagate immune responses and inflammation in the blood vessels, increasing the risk of microthrombosis, which has been documented in lethal and non-lethal COVID-19 cases^1-5^.

One of fundamental mechanisms that EC employs to regulate coagulation is by secreting membrane-associated adhesive glycoprotein von Willebrand factor (vWF) in an elongated secretory organelle, Weibel-Palade body (WPBs)^20-23^. vWF is synthesized in ECs and megakeryocytes and its main function is to directly promote clot formation via capturing platelets and chaperoning clotting factor VIII^22-24^. Current research has confirmed that vWF is not only a plasma glycoprotein known for its clotting role, but also a modulator on inflammatory responses^25-29^. As one of two principal cargos, vWF can determine the assembly of WPB, which contains a variety of other molecules involved in inflammation^21^ and intercellular communication^30^, including angiopoietin-2^27^, IL-8^31^, and eotaxin-3^31^, and intra/extracellular vesicle membrane protein CD63^30^. WPB egress is associated with the secretion of its two major protein constituents, vWF and P-selectin, from EC in response to various stimuli^6,23,32^. P-selectin is a neutrophil and monocyte adhesion molecule important in the initiation of inflammation^18,33^. Collectively, regulating endothelial secretion of vWF from WBP may modulate not only thromboembolic complications, but also inflammation.

Endothelial secretion of vWF in WBP is under tight control^24,34,35^. In general, endothelial WPBs become responsive to exogenous stimuli that increase intracellular Ca^2+^ levels or the second messenger cAMP^36-38^. The effects of cAMP are transduced by two ubiquitously expressed intracellular cAMP receptors, protein kinase A (PKA) and exchange protein directly activated by cAMP (EPAC). EPAC proteins are a family of intracellular sensors for cAMP. In mammals, the EPAC protein family contains two members: EPAC1 and EPAC2^39,40^. Both EPAC isoforms function by responding to increased intracellular cAMP levels in a PKA-independent manner and act on the same immediate downstream effectors, the small G proteins Rap1 and Rap2^40,41^. EPAC1 is the major isoform in ECs, and Rap activation by EPAC1, but not EPAC2, contributes to the effects of cAMP-elevating hormones on endothelial barrier functions^42-44^. Growing evidence has revealed that the cAMP-EPAC signaling axis plays a regulatory role in suppressing inflammation^45,46^. The first identified non-cAMP EPAC1 specific agonist (ESA), I942, was shown to suppress IL-6 signaling and inflammatory gene expression in ECs in response to inflammatory stimuli^46^, suggesting EPAC1 playing an endothelial function-stabilizing role during inflammation^43,47^. In ECs, it has been documented that cAMP provokes the secretion of vWF via the cAMP-PKA pathway^21,35^. We have reported that cAMP-EPAC are involved in hemostasis by driving endothelial luminal surface expressions of tissue plasminogen activator receptor annexin A2, thereby promoting vascular fibrinolysis both *in vivo* and *in vitro*^48^. However, whether cAMP-EPAC is involved in regulating endothelial vWF secretion during inflammation remains to be elucidated. In contrast to the documented EC function-stabilizing effects of the EPACs^43,47^, an *in vitro* study^49^ showed human umbilical vein endothelial cells (HUVECs) increase release of vWF after a short exposure to an EPAC-specific cAMP analog, which can be hydrolyzed by serum esterases^50,51^ and requires to work in starvation media^52^.

In the present study, taking advantage of our *EPAC1*-knockout (KO) mouse model^7^, the EPAC-specific inhibitor (ESI)^53^, and the non-cAMP ESA^52^, we defined the role(s) of EPAC1 in regulation of vWF secretion from ECs in response to inflammatory stimulus both *in vivo* and *in vitro*. We found that the EPAC1 gene deletion elevated the secretion of vWF in endotoxemic mice. Inactivation of EPAC1 potentiated the tumor necrosis factor-α (TNFα)-triggered vWF release. ESA had the capability to reduce the secretion of vWF during inflammation both *in vivo* and *in vitro*. Mechanism studies indicated that activation of host phosphatidylinositol-4,5-bisphosphate 3-kinase (PI3K) / endothelial nitric oxide synthase (eNOS) can attenuate the efficacy of ESI and limit vWFs from secretion during inflammation.

## Materials and Methods

### Ethics Statement

The mouse experiments performed for this study were carried out in accordance with National Institutes of Health, United States Dept. of Agriculture, and UAMS Division of Laboratory Animal Medicine and Institutional Animal Care and Use Committee (IACUC) guidelines. The protocol supporting this study was approved by the UTMB IACUC.

### Endotoxemic mouse models induced by lipopolysaccharide (LPS)

We employed a mouse model of endotoxemia that consisted of intraperitoneal injection of a high dose of *E. coli* LPS (5 mg/kg) (*Escherichia coli* serotype O111:B4; Sigma, MO, USA). For more information, please see Supplemental data.

### Atomic force microscopy (AFM) to measure cell surface expression of target protein

As described previously^54^, the biomechanical properties of P-selectin or CD63 at the cell surface were studied using an AFM system (Flex-AFM, Nanosurf AG, Liestal, Switzerland) that utilized relevant antibody-functionalized AFM probes. For more information, please see Supplemental data.

### Statistics

Statistical significance was determined using Student’s t-test or one-way analysis of variance (ANOVA). Results were regarded as significant if two-tailed P values were < 0.05. All data are expressed as mean ± standard error (SE) of the mean.

## Results

### Inactivation of EPAC1 increases vWF secretion and contributes to the microthrombi progression in LPS-treated *EPAC1*-KO mice

In testing the constitutive plasma levels of vWF, we observed no difference between WT and KO mice (**Figure 1A**). The difference in vWF mRNA expression between these two types of mice was not significant (**Figure 1B**). To further explore the EPAC1 function on inflammation-triggered vWF expression and secretion *in vivo*, WT and KO mice were treated with LPS (5 mg/kg/d, i.p) or PBS for 2 hrs. The results showed that exposure to LPS increased the plasma levels of vWF; compared to the LPS-treated WT counterparts, the plasma levels of vWF were higher in LPS-treated KO mice (*p* < 0.01) (**Figure 1A**). However, the changes of vWF mRNA expressions in lung and liver tissue were not significant (**Figures 1B**). LPS-treated KO mice showed thrombosis, and microthrombi were detected in lungs, brains, and livers of LPS-treated KO mice, but were almost absent in LPS-treated WT mice (**Table 1**) (**Figure S1A**).

**Table 1.**
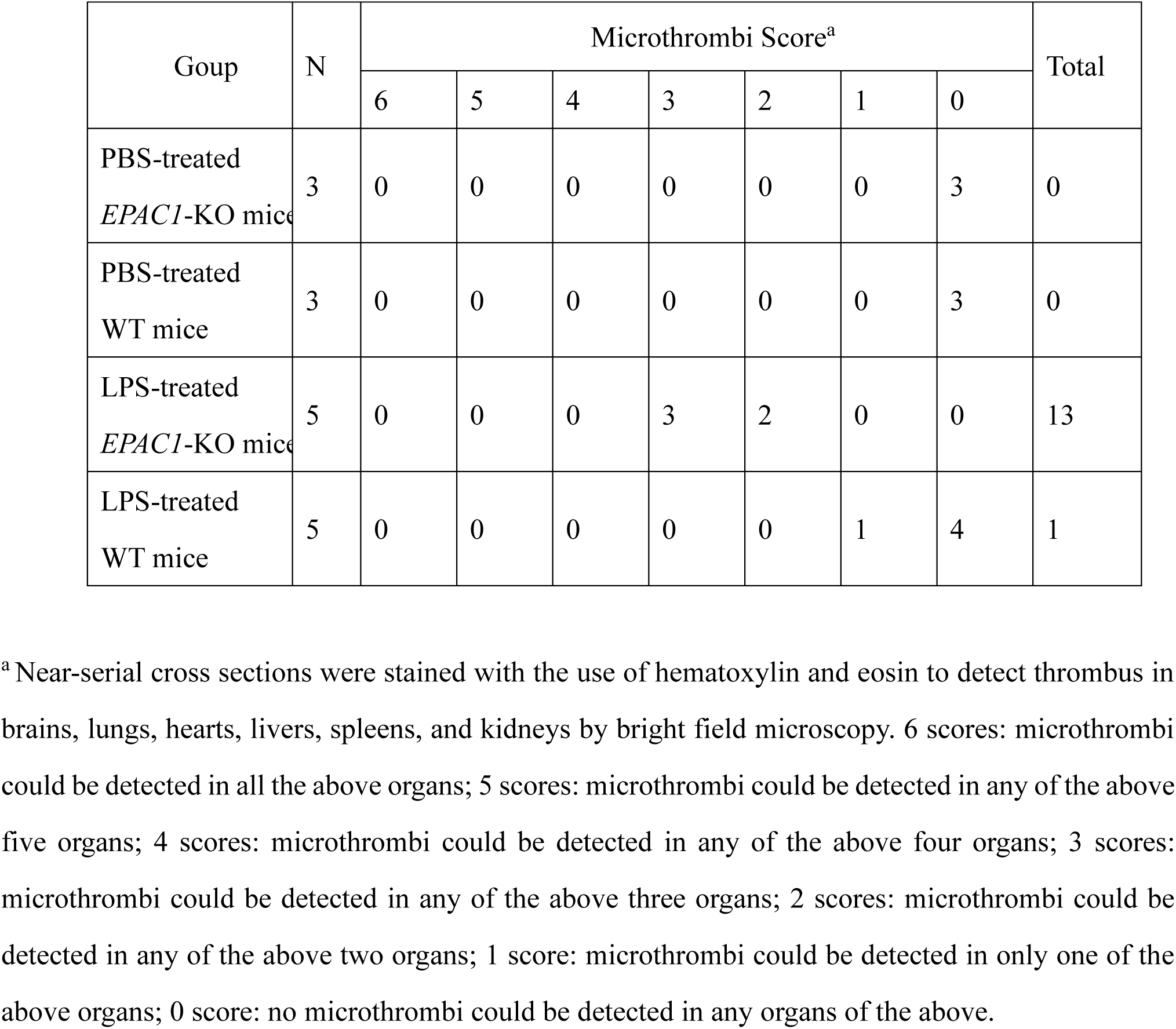
LPS-induced microthrombi detected in mice.

| Goup | N | Microthrombi Score<sup>a</sup> |  |  |  |  |  |  | Total |
| --- | --- | --- | --- | --- | --- | --- | --- | --- | --- |
|  |  | 6 | 5 | 4 | 3 | 2 | 1 | 0 |  |
| PBS-treated<br>*EPAC1*-KO mice | 3 | 0 | 0 | 0 | 0 | 0 | 0 | 3 | 0 |
| PBS-treated<br>WT mice | 3 | 0 | 0 | 0 | 0 | 0 | 0 | 3 | 0 |
| LPS-treated<br>*EPAC1*-KO mice | 5 | 0 | 0 | 0 | 3 | 2 | 0 | 0 | 13 |
| LPS-treated<br>WT mice | 5 | 0 | 0 | 0 | 0 | 0 | 1 | 4 | 1 |

<sup>a</sup> Near-serial cross sections were stained with the use of hematoxylin and eosin to detect thrombus in brains, lungs, hearts, livers, spleens, and kidneys by bright field microscopy. 6 scores: microthrombi could be detected in all the above organs; 5 scores: microthrombi could be detected in any of the above five organs; 4 scores: microthrombi could be detected in any of the above four organs; 3 scores: microthrombi could be detected in any of the above three organs; 2 scores: microthrombi could be detected in any of the above two organs; 1 score: microthrombi could be detected in only one of the above organs; 0 score: no microthrombi could be detected in any organs of the above.

**Fig. 1:**
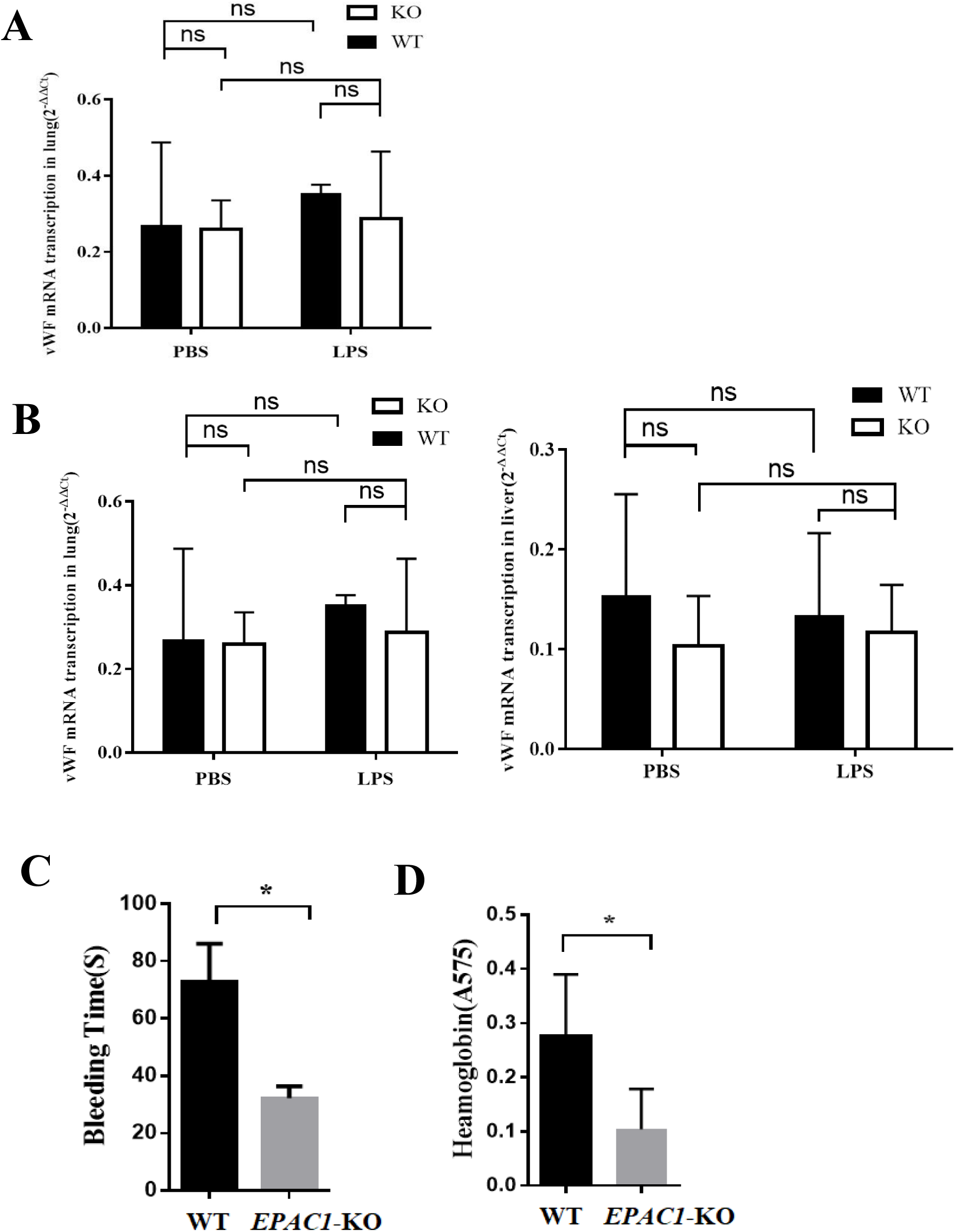
Inactivation of EPAC1 increases vWF secretion. Plasma vWF concentrations and vWF mRNA expression in tissues of WT and *EPAC1*-KO mice treated with or without lipopolysaccharides (LPS). (A)The plasma levels of vWF between WT and KO mice with presence or absence of LPS (5 mg/kg/d, i.p) for 2 hrs, \**P*<0.05, \*\**P*<0.01. n=4 for each group. (B) Reverse transcription-quantitative polymerase chain reaction (qRT-PCR) analysis of vWF mRNA expression in lungs and livers of WT and KO mice treated with or without LPS. Two-Way ANOVA showed that the differences of vWF mRNA expressions were not significant (ns). n=4 for each group (A and B). (C) The bleeding time of WT and *EPAC1*-KO mice treated with LPS. LPS-treated *EPAC1*-KO mice have decreased tail bleeding time, \*\**P* < 0.01. (D) Blood loss quantified as amount of hemoglobin (absorbance at 575 nm) released during the tail bleeding test in LPS -treated WT and *EPAC1*-KO mice, \**P* < 0.05. n=5 for each group (C and D). ns: not significant.

Since vWF performs critical functions in primary hemostasis, a tail bleeding time test was conducted^55^. The bleeding time of LPS-treated KO mice was shorter than that of LPS-treated WT mice (**Figure 1C**). The total blood loss, assessed by its hemoglobin content, corresponded closely with the observed bleeding time of LPS-treated KO and WT mice (**Figure 1D**). Prothrombin time (PT) and activated partial thromboplastin time (aPTT) were also detected. There was no significant difference among the groups (**Figure S1B**).

In conclusion, *EPAC1*-KO mice show a higher level of plasma vWF than WT mice in the state of inflammation, in which microthrombi progression was detected.

### Pharmacological inhibitor of EPAC increases vWF secretion during inflammation in HUVECs

TNFα can regulate vWF expression^56^. We observed that the vWF level was increased in the culture media of HUVECs treated with rTNFα (50 ng/ml) for 4 hrs (*p* < 0.05) (**Figure 2A**). To determine the effect of EPAC1 inhibition on EC secretion of vWF in response to inflammatory mediators, we treated HUVECs with NY173, a novel ESI^53^ to inhibit EPAC1 in HUVECs. NY173 did not affect the viability of HUVECs while the concentration was 1μM to 5μM (**Figure S2A**). NY173(2 μM) had little impact on the vWF secretion from HUVECs (**Figures S2B and S2C**). We found that pretreatment with NY173 (2 μM) for 24 hrs enhanced rTNFα-triggered vWF secretion into media (*p* < 0.01) (**Figure 2A**). Western Blots were used to detect the intracellular vWF protein expression, which was reduced in the rTNFα-treated group (*p* <0.05), and more significantly reduced in the ESI NY173+rTNFα-treated group, vs the rTNF only group (*p* < 0.01) (**Figure 2B**).

**Fig. 2:**
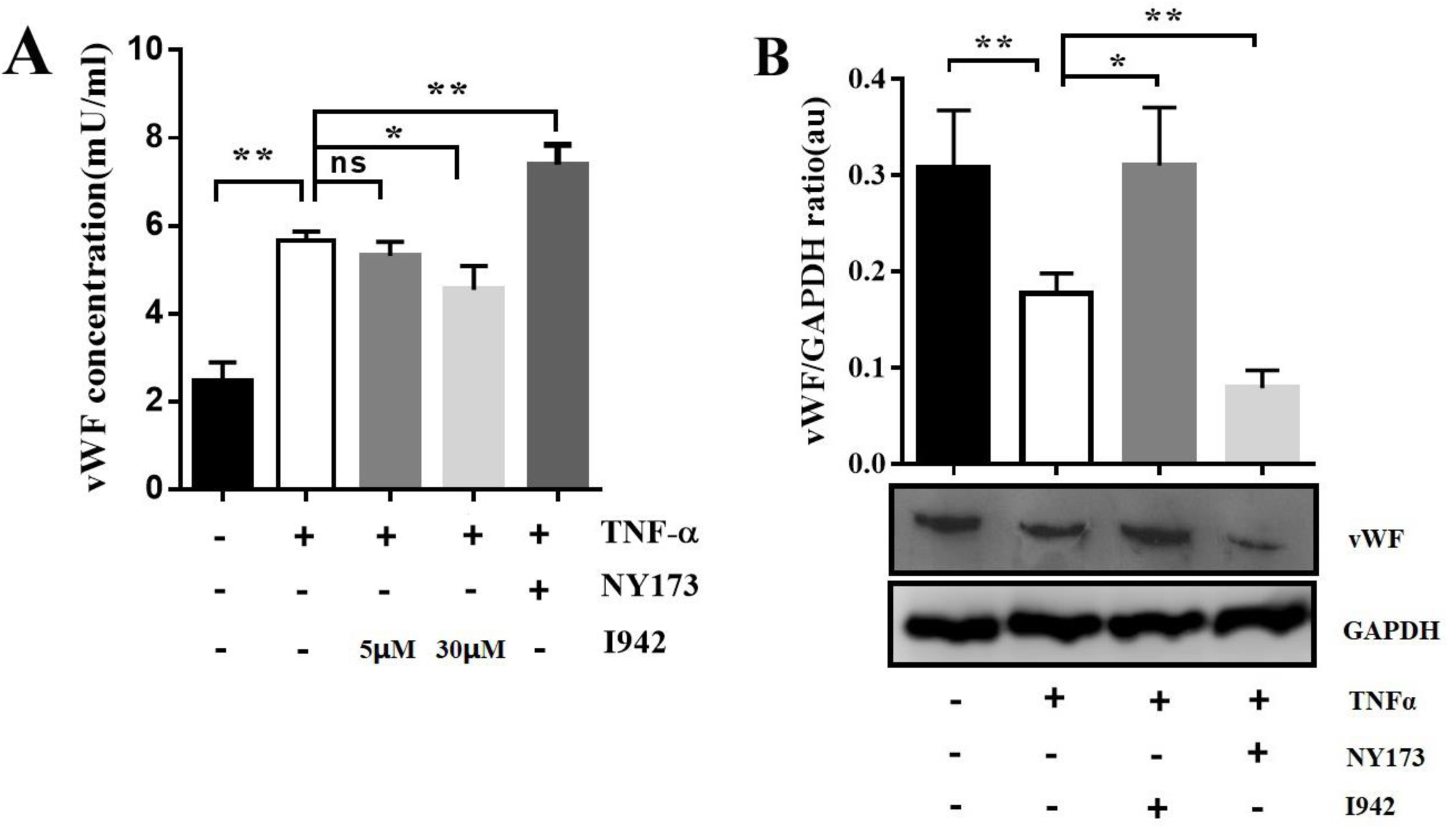
Pharmacological manipulations of EPAC1 modulate inflammation-triggered vWF secretion from HUVECs. (A) vWF concentration in the culture medium of HUVECs. ESI NY173 (2 µM) enhanced the secretion of vWF from rTNFα-treated HUVECs, **P < 0.01. ESA I942 (30 µM) but not ESA I942 (5 µM) has the capability to reduce the secretion of vWF from rTNFα-treated HUVECs, \**P* < 0.05. (B) vWF protein levels in the cell lysis of HUVECs. vWF protein expression reduced in rTNFα-treated HUVECs. vWF protein expression was further decreased in ESI NY173 (2 µM)+ rTNFα group, \*\**P* < 0.01. ESA I942 (30 µM) had the capability to increase vWF expression in rTNFα-treated HUVECs, \**P* < 0.05. n=4 for each group.

### Pharmacological activator of EPAC1 decreases inflammation-triggered vWF secretion from HUVECs

ESA I942, the first identified non-cyclic nucleotide small molecule with agonist properties towards EPAC1 showed very little agonist action towards EPAC2 or protein kinase A (PKA)^57,58^. While the concentration of I942 was among 5μM to 50μM, it had little influence on the viability of HUVECs (**Figure S2A**). I942 (30 μM) had no effect on the vWF secretion from HUVECs (**Figures S2B and S2C**). In this study, HUVECs were pretreated with I942 (30 μM) for 24 hrs before exposure to rTNFα (50 ng/mL) for 4 hrs. ESA I942 (30 μM) down-regulated the secretion of vWF from rTNFα-treated HUVECs (*p* < 0.05) (**Figure 2A**). Compared to rTNFα-treated group, a significantly elevated intracellular vWF expression was observed in I942+rTNFα treated group (**Figure 2B**). Combined with the result that ESI NY173 upregulates rTNFα-induced vWF release, this demonstrates that EPAC1 modulates vWF secretion from HUVECs and the non-cAMP ESA has the capacity to limit vWF from secretion during inflammation.

### EPAC1 modulates the exocytosis of WPBs during inflammation

To illuminate the effect of EPAC1 on the WPB secretion during inflammation, vWF-IF was used to detect intracellular WPBs^59,60^. HUVECs were pre-treated with NY173 (2 μM) or I942 (30 μM) for 24 hrs followed by rTNFα (50 ng/ml) for 4 hrs. IF-labeled intracellular WPBs showed rod or spot shapes in the fluorescent images (arrowheads in **Figure 3A**). Image J analysis showed that the number of WPBs in rTNFα group was decreased significantly, suggesting that WPBs release could be triggered by rTNFα (**Figure 3B**). However, ESA I942 limits the rTNFα-stimulated exocytosis of WPBs, while ESI NY173 enhances such egress (**Figure 3B**).

**Fig. 3:**
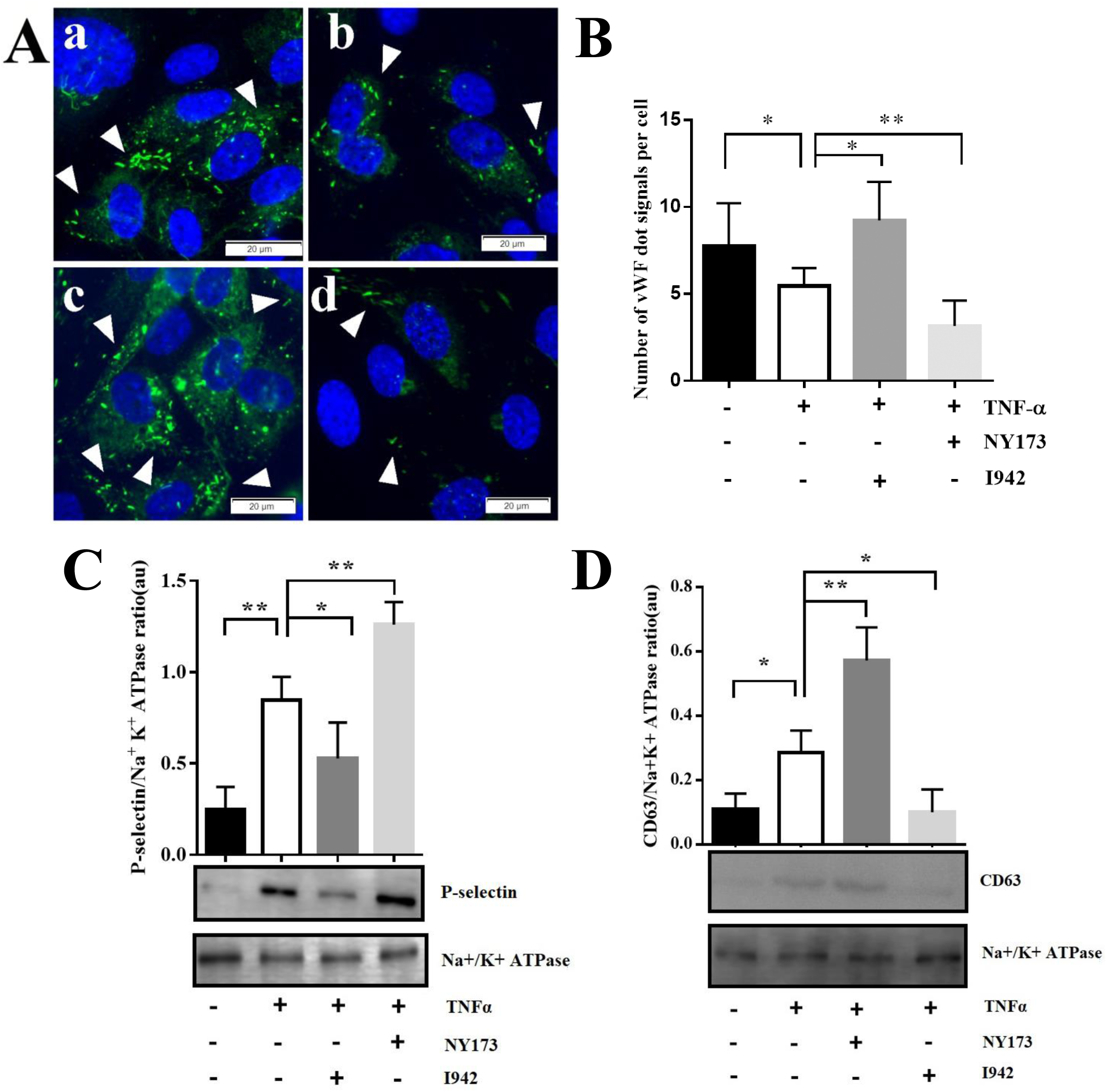
EPAC1 modulates the exocytosis of WPBs during inflammation. (A) Immunofluorescence of vWF in HUVECs. HUVECs were pretreated with NY173 or I942 for 24hrs and treated with rTNFα (50 ng/ml) for 4hrs. The sample was incubated with anti-vWF mouse monoclonal antibody as first antibody, then stained with Alexa Fluor 488-conjugated secondary antibody for vWF labels (green). Nuclei were counter-stained with DAPI (Blue). (a) Vehicle, (b) rTNFα, (c) ESA I942 + rTNFα, (d) ESI NY173 + rTNFα. Scale bars, 20 µm. (B) Quantitative analyses of vWF positive puncta and cell nuclei were performed using Image J software. The results were expressed as dot signals enumerated in each cell. Five microscopic fields were examined for each case. The results were expressed as dot signals enumerated in each cell. n = 6 for each group (A and B). \**P* < 0.05, \*\**P* < 0.01. (C) Western blots were performed to analyze the expression of P-selectin in plasma membrane of HUVECs. (D) Western blot analysis of the expression of CD63 in plasma membrane of HUVECs. Na+-K+-ATPase was used as loading control. n=4 for each group (C and D). \**P* < 0.05, \*\**P* < 0.01.

In addition to vWF and P-selectin, CD63 was detected in WPBs^61^. To further assess whether WPBs exocytosis during inflammation can be modulated by EPAC1, HUVECs were treated with NY173 or I942 for 24 hrs followed by rTNFα (50 ng/ml) for 4 hrs. Increased expressions of P-selectin in plasma membranes from cells treated with rTNFα+NY173 were observed compared with rTNFα groups (**Figure 3C**). The same result was observed in the expression of CD63 (**Figure 3D**). Inverse effects of ESA were showed by observing decreased P-selectin and CD63 expressions in plasma membranes from rTNFα+I942-treated cells (**Figure 3**).

To further validate these results in living cells, we used AFM to quantify the distribution of CD63 and P-selectin on surface of HUVECs. AFM has been used to determine the expression levels of cell surface proteins by measuring the binding affinity of specific protein–protein interactions with nano force spectroscopy^7^. In our study, we measured the specific unbinding force during rupture of the interaction between the antigen (CD63 or P-selectin) expressed at the apical surface of live HUVECs and the antibody-coated AFM cantilever probe. Interactions between antibodies on the AFM cantilever and cell surface antigens caused large adhesion forces, which were quantified by the deflection signal during separation of the cantilever from the cell. The result showed that the adhesion forces in NY173+rTNFα group were stronger than those in single rTNFα group (**Figure 4**), consistent with increased P-selectin and CD63 antigen expression on the apical surface of a single live HUVEC.

**Fig. 4:**
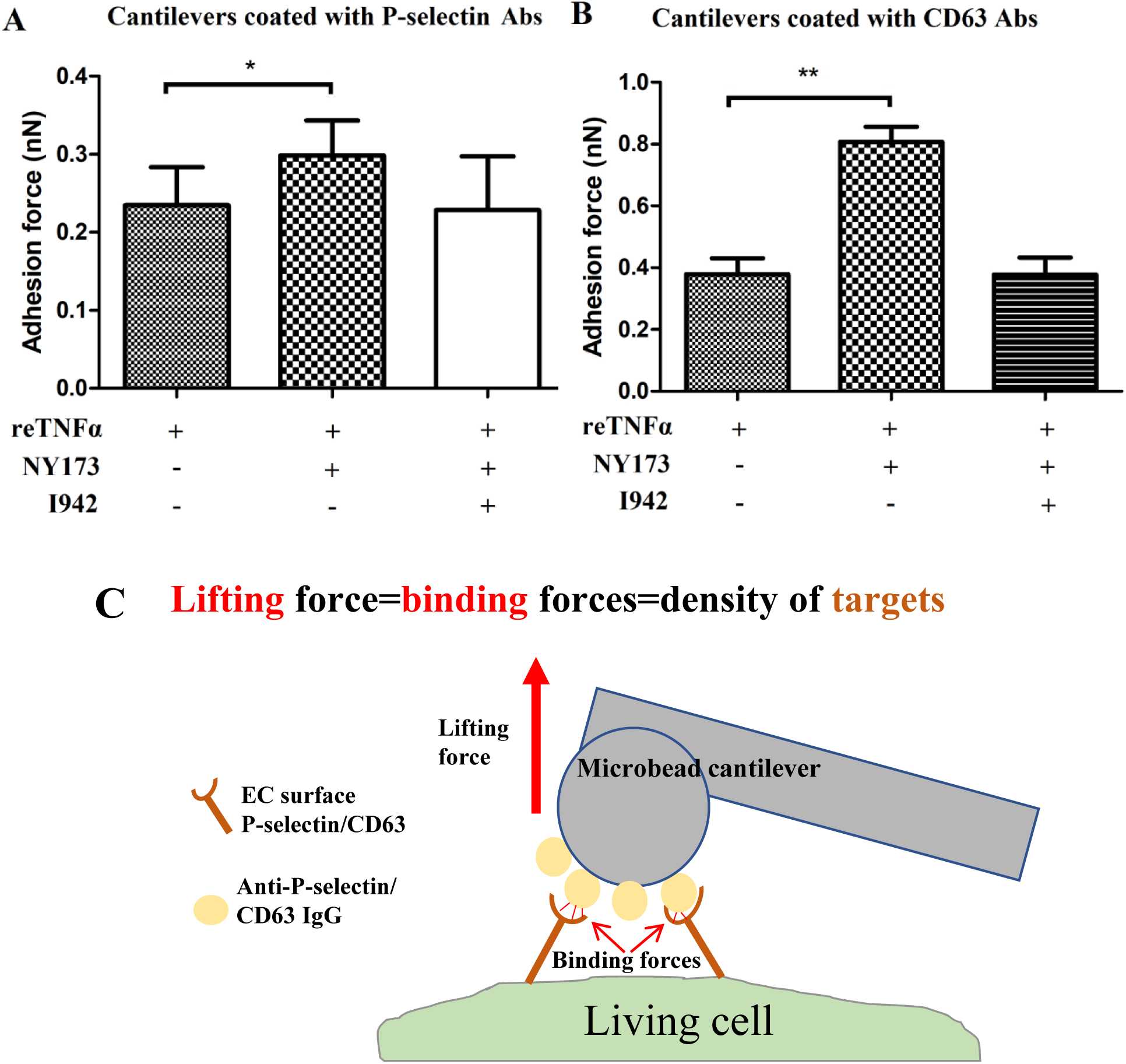
The quantification of P-selectin (A) and CD63 (B) expressions on the living HUVECs surface measured by atomic force microscopy (AFM). Normal mouse IgG-coated colloidal cantilever was employed during calibration of the AFM system before measuring the unbinding forces between P-selectin or CD63 antibody-coated cantilever and the cell (detail see Materials and Methods). (C). The specific unbinding force measurements can be compiled for quantification of total P-selectin or CD63 expressions on certain areas on the living HUVECs surface. With P-selectin or CD63 antibody-coated cantilevers, the adhesion force was stronger in NY173+ TNFα-treated HUVECs, \**P* < 0.05. n=12 for each group.

These data suggest that pharmacological inhibition or activation of EPAC1 affects the expression of P-selectin and CD63 on cell membranes and modulate the inflammatory-triggered exocytosis of WPBs in HUVECs.

### EPAC1 manipulates the spatial proximity between P-selectin and vWF in HUVECs

We next assessed whether P-selectin is only the molecular cargo of WPBs or correlated with vWF closely in EC. Previous studies have shown that P-selectin can bind to D′D3 domains of vWF, which is crucial for P-selectin recruitment^62^. Moreover, D′D3 domains of vWF have already been implicated in vWF storage and multimerization^63,64^. These studies suggest that P-selectin is more than a cargo protein of WPBs. Understanding the complex molecular spatial relationship of protein-protein physical interaction is essential for accomplishing comprehensive knowledge of the functional outcome^65^. In this study, PLA was used to detect P-selectin/vWF spatial proximities and CD63/vWF spatial proximities in HUVECs during inflammation, as we described^65^. Both the P-selectin-vWF proximities and CD63-vWF proximities could be detected in HUVECs. More importantly, pharmacological manipulation of EPAC1 could regulate the P-selectin-vWF proximities (**Figures 5A, B and D**), but not the CD63-vWF proximities (**Figures 5C and E**). PLA signals of P-selectin-vWF in HUVECs were increased in I942 group, while NY173 inducing an opposite effect (**Figure 5D**). The results of PLA suggest that protein-protein proximal interactions exist between vWF and P-selectin or CD63, and that the P-selectin-vWF spatial proximity can be modulated by EPAC1.

**Fig. 5:**
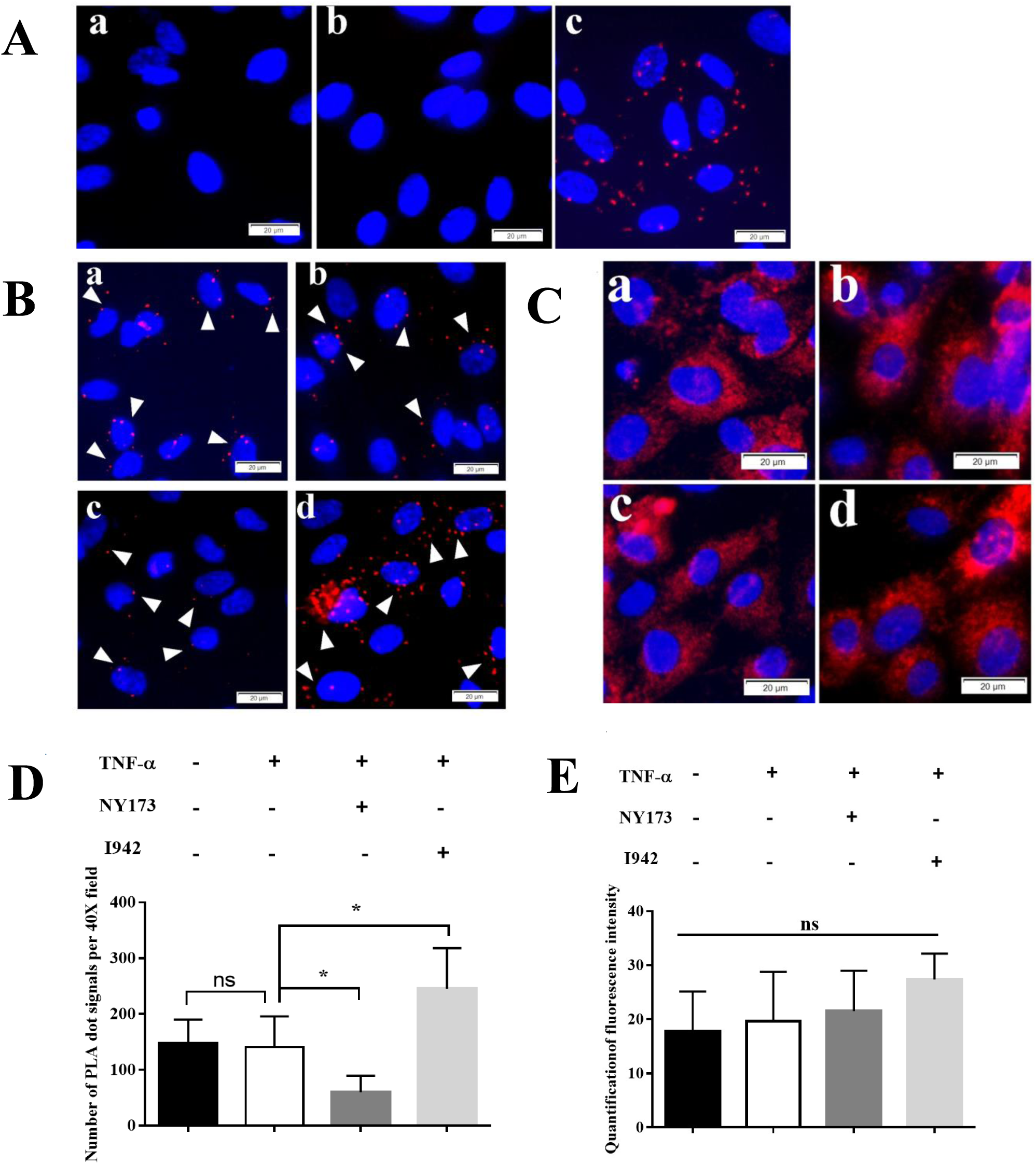
PLA signals of vWF-CD63 and vWF-P-selectin in HUVECs. (A) Control groups. Reagent negative control(a), normal mouse and rabbit IgGs were used as primary Abs. Negative control(b), mouse vWF Abs and Rabbit Rab5 Abs were used as primary Abs. Positive control(c), mouse anti-talin Abs and rabbit anti-α-catenin were used as primary Abs. (B) Representative PLA signals (red) of vWF-P-selectin (arrowheads). (C) Representative PLA signals (red) of vWF-CD63. Vehicle(a), rTNFα(b), ESI NY173 pretreatment+ rTNFα(c), ESA I942 pretreatment+ rTNFα(d). Nuclei were counter-stained with DAPI (Blue). Scale bars indicate 20 µm. (D) Quantitative analysis of vWF-P-selectin PLA signals using Image J software. Five 40× microscopic fields were examined for each case. The results were expressed as dot signals enumerated in each 40× field. n = 3 per group. (E) Quantitative analysis of vWF-CD63 PLA signals using Image J software. Five microscopic fields were examined for each case. The results were expressed as fluorescence intensity per cell. n = 3 for each group.

### EPAC1 regulates TNFα-triggered vWF secretion in a PI3K/eNOS-dependent manner

The nitric oxide (NO) system has a wide range of biological properties in maintaining vascular homeostasis. It is known that the TNFα effect on the NO message is associated with down-regulation of endothelial nitric oxide synthase (eNOS)^66,67^, and that the lack of eNSO likely contributes to the release of vWF^68^. We therefore examined whether rTNFα decreased eNOS expression, and whether it then promoted vWF release from HUVECs. We found that rTNFα reduced the eNOS mRNA but not iNOS mRNA levels in HUVECs (*p* < 0.01) (**Figure 6A and Figure S3**). L-NAME hydrochloride, a NOS inhibitor, showed a non-significant trend to promote rTNFα-induced vWF secretion. Nevertheless, rTNFα-triggered vWF secretion could be markedly reversed by DETA NONOate, an NO donor (**Figure 6B**). These results suggested that TNFα promote vWF secretion through downregulating the expression of eNOS in HUVECs.

**Fig. 6:**
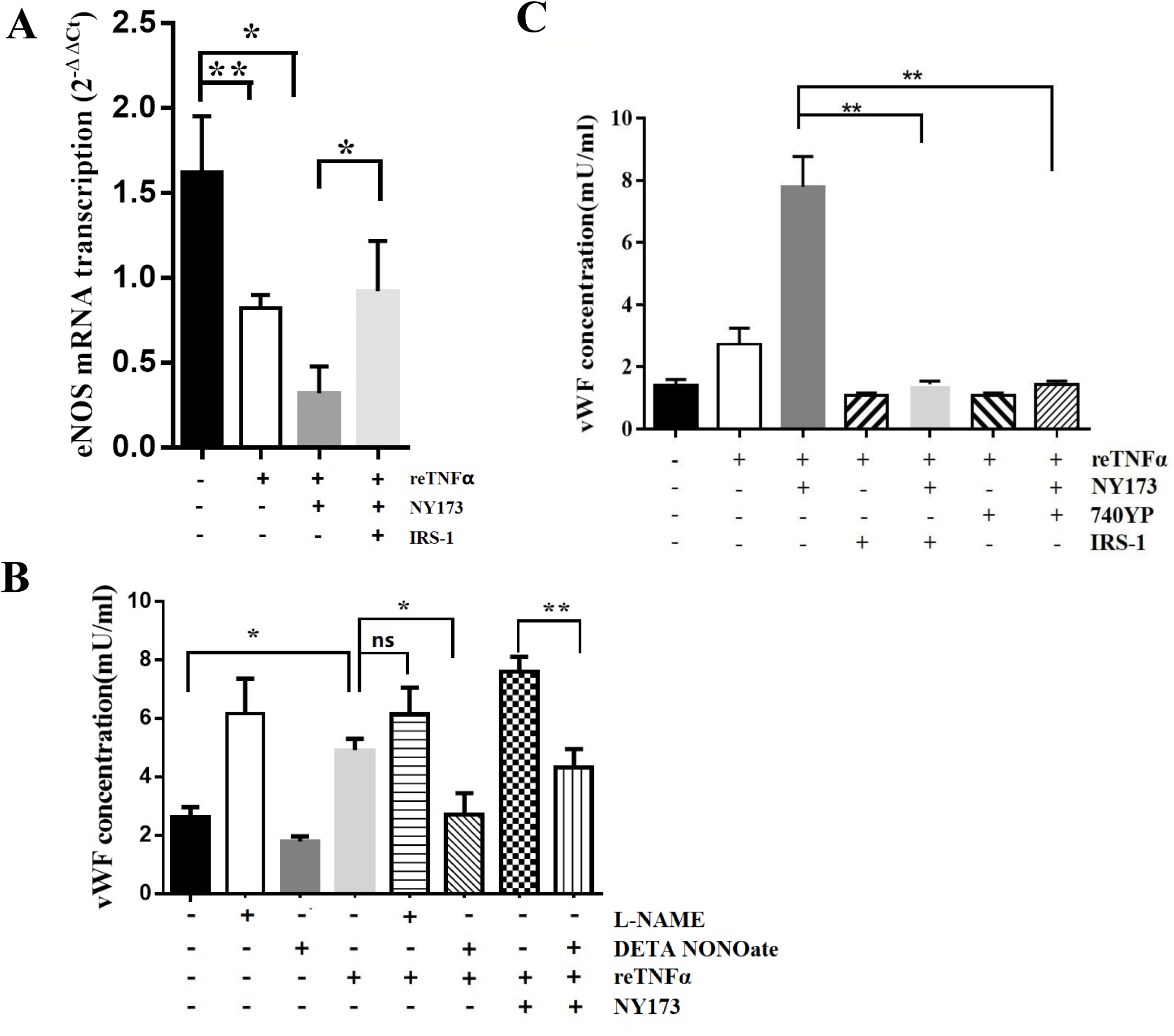
EPAC regulates vWF secretion from HUVECs in a PI3K/eNOS-dependent manner. (A) Reverse transcription-quantitative polymerase chain reaction (RT-qPCR) analysis of eNOS mRNA expression in HUVECs. eNOS mRNA expression decreased in HUVECs treated with rTNFα for 4 hrs; With the present of NY173 for 24 hrs and rTNFα for 4 hrs, the eNOS mRNA expression further declined; PI3K activator IRS-1 reverse the effect of NY173. (B) ELISA assessment of the eNOS role on vWF secretion triggered by rTNFα. DETA NONOate downregulated the rTNFα-induced vWF secretion. The regulation role of EPAC on rTNFα-triggered vWF secretion was eNOS-dependent. (C) ELISA assessment of the PI3K effect on EPAC-regulated vWF secretion from HUVECs. The culture media vWF concentrations were detected in Vehicle-, rTNFα-, rTNFα+NY173-, rTNFα+740YP-, rTNFα+NY173+740YP-, rTNFα+IRS-1-, and re-TNFα+NY173+ IRS-1-pretreated HUVECs. n=3 for each group.

Next, we examined whether the regulatory role of EPAC on rTNFα-triggered vWF secretion was eNOS depended, by comparing pretreatment with NY173 solely, or together with NO donor before challenged with rTNFα. RT-qPCR showed that NY173(2μM) had no effect on the mRNA level of eNOS (**Figure S3A**), but the eNOS mRNA level was further decreased in the NY173+rTNFα group than in rTNFα only group (*p* < 0.05) (**Figure 6A**). However, iNOS did not made such a change (**Figure S3B**). Furthermore, the effect of NY173 on TNF-triggered vWF secretion can be neutralized by NO donor (*p* < 0.01) (**Figure 6B**).

Activation of EPAC leads to a PI3K-dependent PKB activation^69^. EPAC-PI3K-eNOS could be a downstream pathway of adenylyl cyclases^70^. To evaluate the potential effect of EPAC-PI3K-eNOS on the vWF secretion during inflammation, we incubated NY173-pretreated HUVECs with or without PI3K-specific activator IRS-1^71^ before exposure to rTNFα. With the presence of IRS-1, the eNOS mRNA transcription (**Figures 6A**) but not the iNOS transcription (**Figure S3B**) was markedly increased. Furthermore, activation of PI3K in NY173-pretreated HUVECs significantly reduced the rTNFα-triggered vWF secretion compared with the NY173-treated group that did not receive IRS-1 (**Figures 6C**). Another PI3K activator, 740YP (10 μM)^72^, had a similar effect (**Figures 6C**). These results suggest that EPAC1 regulates inflammation-triggered vWF secretion in a PI3K/eNOS-dependent manner.

### ESA I942 reduces the LPS-induced vWF secretion in vivo

To further evaluate the effects of EPAC1 on the vWF secretion during inflammation *in vivo*, WT mice were pretreated with I942 (5 mg/kg/d, i.p.×3) or PBS, followed by LPS injection (5 mg/kg/d, i.p.×1) or PBS. ELISA was performed to detect the plasma vWF concentrations. The results showed that I942 treatment significantly reduced the plasma vWF levels of endotoxemic mice compared with the control group (**Figure 7**), suggesting targeting EPCA1 with ESA can potentially control coagulopathy during inflammation.

**Fig. 7:**
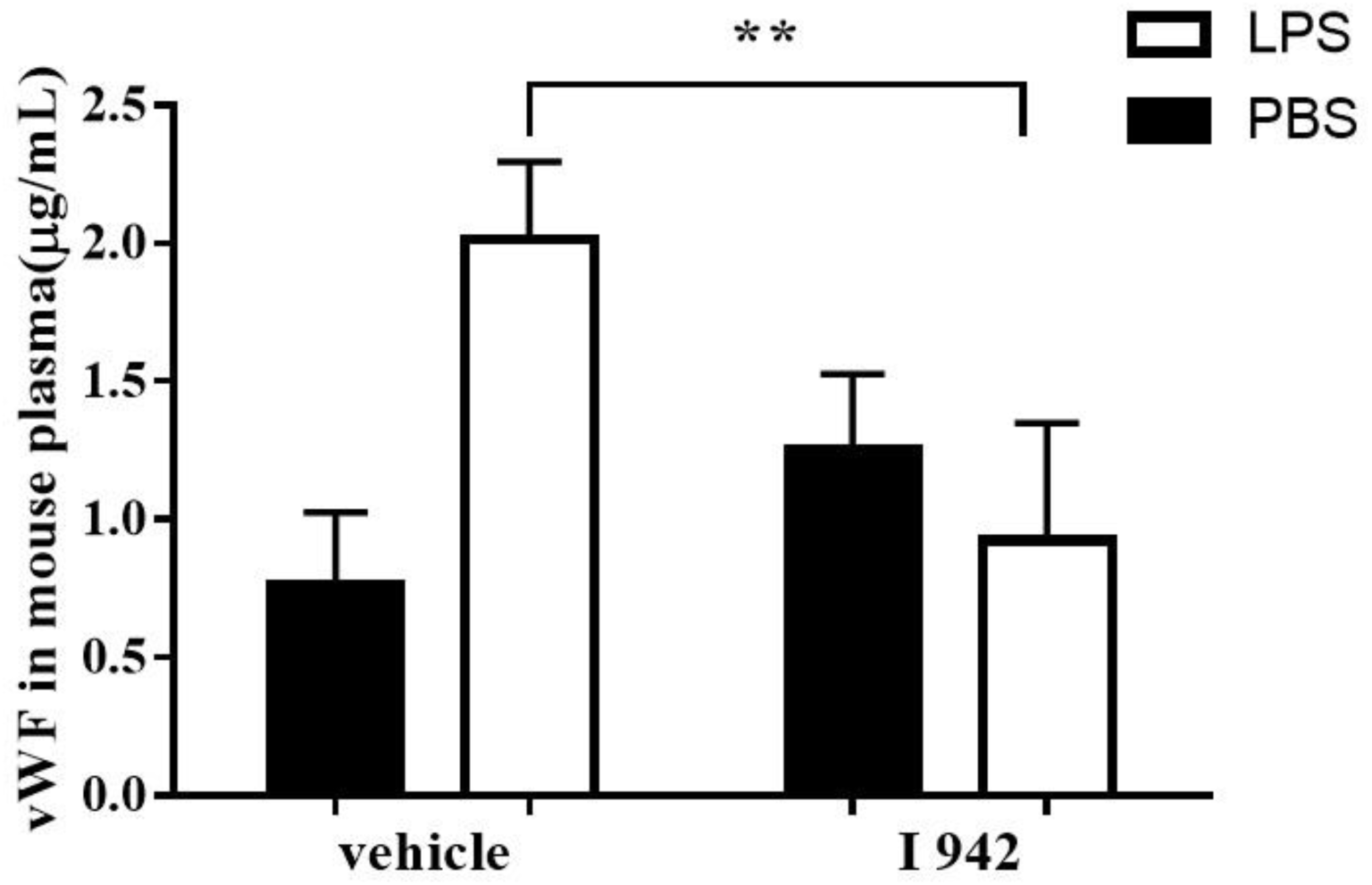
ESA I942 reduces the LPS-induced vWF secretion in WT mice. ELISA was performed to analyze the plasma vWF concentrations in C57BL/6 mice which were treated with I942. Based on unpublished pharmacokinetic and toxicology data from another ongoing project to develop novel ESIs and ESAs, we used I942 at 5mg/kg/day (5 mg/kg/d, i.p.) for three days, then exposed to LPS (5 mg/kg, i.p.) or equal volume of PBS for 2hrs. I942 down regulated the plasma vWF levels in endotoxemic mice, \*\**P*<0.01. n=4 for each group.

## Discussion

Using *in vivo* and *in vitro* models, we demonstrated that EPAC1, as a documented EC function-stabilizing effector, regulated the vWF secretion from ECs triggered by inflammation. Deletion of the *EPAC1* gene or pharmacologically inactivation of EPAC1 significantly facilitated inflammation-induced vWF secretion. On the contrary, activation of EPAC1 by ESA attenuated the vWF secretion in ECs under inflammation. ESI increased P-selectin expression in cell plasma membrane and affected the protein space proximity of P-selectin-vWF. Activation of PI3K or increased NO expression in NY173-treated HUVECs significantly reduced the efficacy of NY173 in promoting inflammation-triggered vWF secretion. Furthermore, by utilizing the endotoxemic mouse model, we observed that ESA (I942) reduced the inflammation-induced vWF secretion *in vivo*. These *in vivo* and *in vitro* data unravel a novel regulatory role for EPAC1 in vWF secretion from ECs. Importantly, the cAMP-EPAC signaling axis can be exploited as a potential target for the development of compounds to control vWF secretion triggered by inflammation.

The hallmark of acute and chronic inflammation is the widespread activation of ECs which provokes excessive vWF secretion from the EC storage pool by WPB exocytosis^73^. vWF is secreted via three pathways. They are regulated secretion, basal secretion, and constitutive secretion^23,34^. The first two types of vWF-secretion pathways occur from WPBs and deliver highly multimerized vWF, but the third pathway releases vWF that has not been packaged into WPBs and, thus, has not undergone high levels of multimerisation^38^. In contrast to continuously secreting vWF from basal pathway, the release of vWF from the regulated pathway occurs only following stimulation of ECs with an appropriate agonist, providing the endothelium with the means to react to its microenvironment by finely tuning the rate of release. Thus, controlling WPB exocytosis from ECs is an important way to regulate the concentration of vWF in plasma under inflammatory stress^73,74^.

It has been proved that treatment of HUVECs with I942, which represents an effective tool to probe the function of cellular EPAC1, leads to alterations in the expression of a wide variety of genes associated with vascular function. In particular, I942 suppresses the expression of the proinflammatory adhesion molecules in HUVECs^75^. Upon an inflammatory trigger, ECs are activated, prompting the massive release of WPBs that contain vWF and P-selectin^76^. We observed rTNFα-induced vWF secretion was increased in ESI pretreatment group, while ESA had the opposite effect. In addition to these *in vitro* results, *EPAC*-KO endoxemic mice had higher level of plasma vWF concentration than WT endoxemic mice. The results confirm that EPAC1 plays an important role in regulating inflammation-triggered vWF secretion. In contrast to our study, van Hooren, et al. reported that Me-cAMP-AM, an EPAC-specific cAMP analog, promotes release of vWF in HUVECs^49^. This discrepancy could be due to the different reagents and experimental conditions used in the *in vitro* experiments. In van Hooren’s study, HUVECs were incubated with serum-free medium and supplemented with 1 μM Me-cAMP-AM for 60 mins. Since cAMP analogs are hydrolyzed by serum esterases^50,51^ and requires to work in starvation media^52^, which restricts applications using primary ECs^47,77,78^. Moreover, evidence has been provided that most cAMP and cGMP analogs have multiple targets, including some EPAC specific analogs^77,78^. In our study, we used the non-cyclic nucleotide ESA I942, to treat HUVECs for 24 hrs followed by rTNFα (50 ng/mL) for 4 hrs with complete medium. We observed not only that ESA could inhibit vWF secretion triggered by inflammation, but also that ESI increased inflammation-triggered vWF secretion, in accordance with our observation during *in vivo* study using WT and KO mice.

To our knowledge, P-selectin, as a critical component of WPB, is anchored to the cell surface with the exocytosis of WPB following inflammatory stimulation of endothelial cells, and then provides binding sites for P-selectin ligands on the surface of circulating leukocytes^79,80^. In the present study, P-selectin expression in the plasma membrane was elevated in rTNFα-treated HUVECs with ESI pretreatment, while the expressions were attenuated in that group with ESA pretreatment. The expressions of CD63, an essential cofactor to leukocyte recruitment by endothelial P-selectin^30^, were consistent with the change of P-selectin expressions. To our knowledge, this is the first report that EPAC regulates the expressions of P-selectin and CD63 on the cell membrane following inflammatory stimulation. Our results confirm the well-documented EC function-stabilizing effects of the EPACs from another perspective. Moreover, P-selectin, as a plasma marker of endothelial damage/dysfunction, is more than a cargo protein of the WPB. It has been reported that P-selectin binds to the D-D3 domains of VWF, which is not only crucial for P-selectin recruitment, but also be implicated in VWF storage^62,63^. In our study, PLA was used to detect the spatial proximity between P-selectin and vWF, showing that pharmacological manipulation of EPAC1 can regulate P-selectin-vWF spatial proximities. The domains of vWF binding with P-selectin is critical to the storage of vWF. Whether EPAC1 can manipulate the secretion of vWF by regulating the binding of P-selectin and vWF remains to be determined.

Under normal physiological conditions, ECs help prevent inflammation and inhibit clotting partly through the continuous production of nitric oxide^81,82^. Viral infections, such as severe acute respiratory syndrome coronavirus-2 (SARS-CoV-2), cause inflammatory cytokines release, EC injury, and NO production decline significantly^83^. TNFα can downregulate the eNOS expression in ECs^66^. Moreover, eNOS reduction leads to vWF release^68^. In this study, we also observed that rTNFα (50 ng/mL) incubating HUVECs for 4 hrs could potently decrease the eNOS mRNA expression. Using an NOS inhibitor or NO donor to regulate NO expression in HUVECs can manipulate the vWF secretion triggered by TNFα. It has been reported that EPAC activation can enhance the eNOS activity in ECs^84^. Likewise, we observed that ESI NY173 further decreased the TNFα-induced eNOS mRNA reduction. The NO donor, in turn, counteracted vWF secretion from HUVECs under the challenge of NY173 and TNFα. PI3Ks have been linked to an extraordinarily diverse group of cellular functions, including inflammation and coagulation^85-87^. An earlier study has shown that PI3K is critical in mediating the prothrombotic potential of ECs^88^. *In vivo* studies have shown that inhibition of PI3K strongly enhances the activation of EC, upregulates the LPS-induced coagulation and inflammation, and reduces the survival time of mice^87^. We incubated NY173-pretreated HUVECs with the PI3K-specific activators IRS-1 before challenging with rTNF-α. We found IRS-1 could dramatically reverse the effect of NY173 on inhibiting the eNOS expression and releasing of vWF. Activation of PI3K in NY173-pretreated ECs significantly reduced the TNFα-induced inhibition of eNOS expression and vWF secretion, compared with the NY173-pretreated group that did not receive IRS-1. In support of the conclusion that EPAC mediates eNOS expression through the PI3K-Akt pathway from Namkoong et al^70^, we hypothesize that EPAC may regulate the rTNFα-trigger vWF release in a PI3K-eNOS dependent manner. Van Hoooren et al has reported that PI3K inhibitor LY294002 reduced epinephrine-induced release of vWF^89^. Comparing this study with ours, we found considerable differences in reagents and experimental conditions and the time-points observed in these two experiments were different. The incubation time of epinephrine and LY294002 in Van Hooren’s study was ranged from 10 to 50 min, while our incubation time of PI3K activators was 24 hrs followed with 4 hrs of rTNFα-treatment time. In addition, epinephrine was used in Van Hooren’s study to activate the vWF secretion. As we know, epinephrine not only stimulates vWF release through cAMP/EPAC pathway^37,49^, the cAMP/PKA pathway is another pathway that epinephrine regulates and affects vWF secretion^90-92^. EPAC and PKA might mediate the opposing effects on PKB regulation. Activation of EPAC leads to a PI3K-dependent PKB activation, while stimulation of PKA may inhibit PKB activity^69^.

Despite significant progress in the development of cAMP analogues as EPAC agonists^93^, the EPAC isoform selectivity of these cAMP analogues and their potential off-target effects on other molecules, such as cAMP phosphodiesterase, remain challenging^77,78^. ESA I942, the first identified non-cyclic nucleotide small molecule with agonist properties towards EPAC1, and very little agonist action towards EPAC2 or PKA^57,58^, has the potential to suppress proinflammatory cytokine signaling. This reduces the risk of side effects associated with general cAMP-elevating agents that activate multiple response pathways in HUVECs^58^. In this study, it displayed a capability to exert its pharmacological effects in complete EC culture medium and it downregulated inflammation-triggered vWF secretion in HUVECs. Moreover, this report about *in vivo* application of I942 shows activation of EPAC1 *in vivo* significantly reduced the plasma vWF concentration in LPS-induced endotoxemic mice. Additional work is needed to further explore the possibility whether I942 can act as a novel therapeutic to regulate hemostasis.

Taken together, this study combined an *in vitro* primary human endothelial system with an *in vivo* mouse model to reveal the role of EPAC in inflammation-triggered vWF secretion. In vascular ECs, EPAC1 regulates inflammation-triggered vWF releasing in the PI3K/eNOS-dependent manner (**Figure S4**). ESA I942 has the capacity to limit vWFs from secretion during inflammation *in vivo*. These data shed light on potential development of new strategies to controlling the risk of thrombosis during inflammation.

## Acknowledgment

We gratefully acknowledge Dr. Katherin Hajjar and Dr. Zongdi Fen for the scientific advice during this project. We thank Drs. Sarita Paulino, Juquan Song, Amina EI ayadi, Jayson Jay, and Changcheng Zhou; summer students Jianni Bei and Yixuan Zhou for technical supports. We gratefully acknowledge Dr. Kimberly Schuenke for her reviews and editing the manuscript. This work was supported by NIH grants R01AI121012 (B.G.), R21AI137785 (B.G.), R21AI154211(B.G.), R03AI142406 (T.S. and B.G.), R21AI144328 (T.S. and B.G.), and R01AI132674 (L.S.). The funders had no role in the study design, data collection and analysis, decision to publish, or preparation of the manuscript.

## Authorship Contributions

J.X., B.Z., Z.S., Y.L., Q.C., and P.W. performed experiments. J.X., B.Z., Z.S., Y.L., A.G., and B.G. analyzed data. J.X., Z.S., A.B., Y.J., A.G., S.L.R., M.L., T.S., and B.G. designed the study. T.K., J.Z., M.L., and T.S. provided reagents. J.X., B.Z., Z.S., T.R.S., A.B., L.S., T.K., S.L.R., J.Z., M.L., and B.G. wrote the manuscript.

