## supplemental material for "EPAC regulates von Willebrand factor secretion from endothelial cells in a PI3K/eNOS-dependent manner during inflammation"

**Materials and Methods**

**Cell culture**

HUVECs (Cell Applications, Atlanta, GA) were cultured in Endothelial Cell Growth Medium (Cell Applications) with humidity in 5% $CO_2$ at 37°C. The media was changed every 48 hrs until cells reached 90% confluence. Cells passaged 2-5 times were used for all experiments. To stimulate HUVECs, the cells were cultured in media supplemented with 50 ng/mL recombinant TNFα (rTNFα, Sigma, St. Louis, MO) for 4 hrs[1]. Cells were then fixed by 4% paraformaldehyde solution in phosphate-buffered saline (PBS) for 30 min at room temperature (RT) before further processing.

**EPAC inhibitor and activator**

EPAC-specific inhibitor (NY173) and activator (I942) were synthesized in Dr. Jia Zhou's laboratory at UTMB[2,3]. To determine the effects of NY173 and I942 on HUVEC viability *in vitro*, cytotoxicity was determined using the MTT Cell Proliferation Assay kit from Cayman (MI, USA) after culturing HUVECs with different concentrations of (1, 2, 3, 5, µmol/L) of NY173 and (5, 10, 30, 50 µmol/L) of I942, respectively, for 24 h. Mock-treated HUVECs served as controls. H2O2 (200 µmol/L)-treated HUVECs served as positive control. The absorbance at 490 nm was recorded using a BioTek ELx808 plate reader (BioTek, Winooski, VT, USA). Results are presented as mean ± standard error of the mean of three independent experiments; each experiment was conducted in triplicate.

**Special reagents**

The PI3K-specific activator 740YP and the NOS inhibitor L-NAME hydrochloride (L-NAME) were purchased from Tocris Bioscience (Bristol, UK); the insulin receptor substrate (IRS-1 $Tyr^{608}$) was purchased from Enzo Life Sciences (NY, US); Nitric Oxide donor Diethyleneriamine NONOate (DETA NONOate) was from Cayman Chemical (MI, US). For *in vitro* studies, the final concentration of NY173, I942, 740YP, IRS-1 $Tyr^{608}$, DETA NONOate and L-NAME in the HUVECs culture medium were 2 µmol/L, 30 µmol/L, 20 µmol/L, 50 µg/L, 100 µmol/L, and 100 µmol/L, respectively. Medium with 0.01% dimethyl sulfoxide (vol/vol) was used as the vehicle control.

**Plasma membrane protein isolation and cell fractionation**.

Plasma membrane protein isolation and cell fractionation were performed using the Minute™

Plasma Membrane Protein Isolation and Cell Fractionation Kit (Invent Biotechnologies, Eden Prairie, MN), as we described[4,5]. The resulting fractions were resuspended in RIPA buffer for immunoblotting to detect the expressions of P-selectin and CD63 in plasma membrane. Purity of the plasma membrane fractions was verified by immunoblotting with antibody raised against membrane-specific (Na+, K+ ATPase) proteins[5].

**Proximity Ligation Assay (PLA)**

The spatial proximities of P-selectin-vWF or CD63-vWF was determined using the Proximity Ligation Assay, as we described recently[6]. Information for special reagents is as following: Duolink® In Situ PLA® Probe Anti-Rabbit PLUS, Affinity purified Donkey anti-Rabbit IgG(H+L) (Millipore Sigma, DUO92002-30RXN), Duolink® In Situ PLA® Probe Anti-Mouse MINUS, Affinity purified Donkey anti-Mouse IgG(H+L) (Millipore Sigma, DUO92004-30RXN), Duolink® In Situ Detection Reagents Red (Millipore Sigma, DUO92008-30RXN), Duolink® In Situ Mounting Medium with DAPI (Millipore Sigma, DUO82040-5ML), Duolink® Blocking Solution (Millipore Sigma, DUO82007-4ML), Duolink® Antibody Diluent (Millipore Sigma, DUO82008-2.5ML), Duolink® In Situ Washing Buffer A (Millipore Sigma, DUO82046-1EA), Duolink® In Situ Washing Buffer B (Millipore Sigma, DUO82048-1EA). Briefly, the samples were treated with 0.25% TritonX-100 (diluted in PBS) for 10 min at RT, and then blocked with Duolink® blocking solution in shaker incubator at 37°C for 1 hr. After incubation with designed primary antibodies, the samples were incubated with MINUS probe and PLUS probe. 1×Ligase (1unit) added in 1×Ligation (39units) was added to the sample before Duolink amplification. We used the spatial proximities of talin/α-catenin as a signal positive control and vWF-Rab5 as a negative control, respectively, in HUVECs for PLA[6]. For a reagent negative control, mouse normal IgG and rabbit normal IgG were used as the primary antibodies. All samples were analyzed with an Olympus BX51 fluorescence microscope.

**Endotoxemic mouse models induced by lipopolysaccharide (LPS)**

In this study, C57BL/6 *EPAC1* null (KO) mice were derived as described previously[7]. All mice were purpose-bred at UTMB and used between 8 and 12-week-old for all experiments. KO mice were viable, fertile, and without overt abnormalities. We employed a mouse model of endotoxemia that consisted of intraperitoneal injection of a high dose of *E. coli* LPS (5 mg/kg) (*Escherichia coli* serotype O111:B4; Sigma, MO, USA)[8-10]. LPS, which is derived from Gram-negative bacteria, binds the LPS binding protein. In response to LPS, monocytes produce large quantities of proinflammatory cytokines including TNFα[11]. To measure vWF concentration in plasma, blood samples were collected 2 hrs after LPS treatment. After 24 hrs of LPS treatment, organs were dissected after euthanasia and perfusion via right ventricle[12].

**Bleeding time, hemoglobin measurement, prothrombin time (PT), and activated partial thromboplastin time (aPTT) test**

WT and *EPAC1*-KO mice were anesthetized with isoflurane. The distal 5 mm of the tail was clipped off with a scalpel. The bleeding tail was placed in 37°C in PBS. The time until bleeding cessation was recorded as the bleeding time[13]. Bleeding was also indirectly assessed by measuring the resulting hemoglobin content in the PBS solution. After centrifugation, blood cells were precipitated, and 1 mL RIPA buffer was added for cell lysis. Absorbance was read at 575 nm in a BioTek™ ELx808™ Absorbance Microplate Reader.

Plasma samples from citrated blood collected from WT and *KO* mice were used to determine prothrombin time (PT) and activated partial thromboplastin time (aPTT). Thromboplastin-D and aPTT-XL (Ellagic Acid Activator) with CaCl2 were purchased from Thermo Fisher Scientific (NH, US). The PT and aPTT were determined following the manufacturer's instructions, as described previously[4].

**Histopathology, Immunofluorescence (IF), and Enzyme-linked immunosorbent assay (ELISA)**

Complete necropsies were performed on all experimentally treated and control mice. Samples of brain, liver and lung tissues were fixed in a 4% (vol/vol) neutral buffered solution of formaldehyde, embedded in paraffin, sectioned at 5-μm thickness, and processed by staining with hematoxylin and eosin for evaluation of histopathology. IF of vWF were performed in HUVECs. The sample was incubated with anti-vWF mouse monoclonal antibody (1:500) (Thermo Fisher Scientific). After 2 hrs, the samples were incubated with Alexa Fluor 488-conjugated goat anti mouse IgG (1:2000) or Alexa Fluor 594-conjugated goat anti rabbit IgG (1:1000) for 1 hr. A mouse monoclonal IgG (Thermo Fisher Scientific, NH, US) was served as a negative control. Nuclei were counterstained with DAPI. Fluorescent images were taken and analyzed using Olympus BX51-microscope, as described previously[6].

Based on the IF images, quantitative analyses of WPB were performed using Image J software (National Institutes of Health, USA)[14]. Briefly, image thresholding was processed to create binary pictures revealing vWF positive puncta or cell nuclei, and then the size range of the puncta using the wand tool were determined. After the parameters were set, we used particle analyzer to quantify the number of the WPBs puncta and cell nuclei. The number of WPBs normalization was done by dividing the number of WPBs by the number of cell nuclei in the same view.

Plasma samples from citrated blood collected from WT and EPAC1-KO mice were used for ELISA to detect mouse vWF (ABclonal, MA, US). The vWF concentrations in culture medium from HUVECs were detected by human VWF ELISA Kit (Assaypro, MO, US). Standard curves were created by serial dilution of standard proteins provided in the kits. All ELISA plates were read at 450 nm, according to the manufacturer's directions.

**Real-time quantitative polymerase chain reaction (qRT-PCR)**

Total RNA was prepared from tissue or cell samples using TRIzol (Life Technologies, CA, USA). Total RNA was quantitated using NANO Drop 2000. Complementary DNA was prepared with iScript Reverse Transcription Supermix for qRT-PCR (Bio-Rad, CA, USA), and qRT-PCR was carried out with primers for mouse vWF (Fwd: 5'-AAC AGA CGA TGG TGG ACT CAGC-3' and Rev: 5'-CGA TGG ACT CAC AGG AGC AAGT-3'）, human eNOS (Fwd: 5'-GGC TCA GTT ACT GTC TAA GTG TTA GAA-3' and Rev: 5'-CCC TGG AGT CTT GTG TAG GAT AT-3'), human iNOS (Fwd: 5'-CCT GGC AGC CAT TTC AGA GGA G-3' and Rev: 5'-CCA GCC TCA AGT CTT ATT TCC TCA A-3'), and iTaq Universal SYBR Green PCR Supermix (Bio-Rad, CA, USA). We normalized both mouse and human data by using housekeeping genes, actin and glyceraldehyde-3-phosphate dehydrogenase (GAPDH), as references.

**Atomic force microscopy (AFM) to measure cell surface expression of target protein**

AFM is an advanced tool for studying biomechanical properties and has been used to determine the expression levels of cell surface proteins by measuring the binding affinity of specific protein–protein interactions, including antigen–antibody and receptor–ligand, with nano force spectroscopy[15,16]. In our study, anti-P-selectin or anti-CD63 antibodies were immobilized on polystyrene spheres attached to a colloidal cantilever. We measured the specific unbinding force during rupture of the interaction between the antigen (P-selectin or CD63) expressed in designated fields at the apical surface of living HUVECs and the antibody-coated AFM cantilever probe. Interactions between antibodies on the AFM cantilever and cell surface antigens cause large adhesion forces, which are quantified by the deflection signal during separation of the cantilever from the cell. By tracking the cantilever deflection and retraction cycle, the binding, stretching, and rupture of antibody–antigen complexes can be monitored in terms of the adhesive force changes on the cantilever over the distance traveled by the cantilever. We calculated the work required to break all interactive bonds between the cantilever and the EC, reflecting the quantity of antigen expression on the surface[15,16]. As described previously[16], the biomechanical properties of P-selectin or CD63 at the cell surface were studied using an AFM system (Flex-AFM, Nanosurf AG, Liestal, Switzerland) that utilized relevant antibody-functionalized AFM probes. Colloidal cantilevers with a 5 μm polystyrene bead were used (SHOCON-G-PS, Applied NanoStructures, Mountain View, CA) to measure surface protein interaction forces[15,16]. The cantilevers were functionalized by incubation with anti-P-selectin mAb (Thermo Fisher Scientific) or anti-CD63 mAb (Thermo Fisher Scientific) at 100 μg/ml in 0.1 M NaHCO3 (pH 8.6) overnight at 4°C. Normal mouse IgG was used as negative control during calibration. Unbound proteins were rinsed off using PBS. The exposed surface of the bead was blocked by bovine serum albumin (Sigma, MO, USA) at 500 μg/ml in PBS. AFM imaging and measurements were generally taken within 1 hr after blocking. The spring constant of the cantilever was calibrated using the Sader method in air[17]. The cantilever spring constant varied between 0.10-0.15 N/m. Force spectroscopy was done in static force mode operating on 25 $\mu m^2$ areas on a living cell surface. The functionalized cantilever was manipulated into contact with the surface of a confluent monolayer of HUVECs. The maximum compression force was set to 150 pN. The contact time was kept constant at 500 msec before the cantilever was retracted at a constant pulling speed of 1 μm/s to measure the force-extension curve. We scanned five cells per group, each with a different cantilever.

**Supplemental Figures:**

Figure S1: **The changes of coagulative state of LPS-treated *EPAC1*-KO mice.** (A) The formation of microthrombi in micro vessels of LPS-treated *EPAC1*-KO mice. WT and *EPAC1*-KO mice were treated with LPS (5 mg/kg/d, i.p.×1). After 24 hrs, organs were dissected after mice euthanasia and whole animal perfusion. Organs (liver, spleen, lung, brain, heart, and kidney) were immersion-fixed in 10% buffered formalin overnight. Microthrombi were detected in LPS-treated *EPAC1*-KO mice (a, b, c), but not in LPS-treated WT mice (d, e, f). (a, d) liver, (b, e) lung, (c, f) brain. Scale bars indicate 50 μm. (B) Prothrombin time (PT) and activated partial thromboplastin time (aPTT) of LPS-treated WT and *EPAC1*-KO mice. The differences of PT and aPTT between these two groups were not significant.

Figure S2: **The effect of NY173 and I942 on HUVECs viability and vWF secretion.** (A) The viability of HUVECs was detected by MTT assay. HUVECs were incubated with different concentrations of (1, 2, 3, 5 μmol/L) NY173 and (5, 10, 30, 50 μmol/L) I942 separately for 24 h. 0.1% DMSO-treated HUVECs served as controls. 200 μmol/L H2O2-treated HUVECs served as positive control. n=3 for each group. (B)The effect of NY173 and I942 on vWF secretion. HUVECs were incubated with 2μmol/L NY173 or 30 μmol/L I942 for 24 hrs. The vWF concentrations in culture medium from HUVECs were detected by Human VWF ELISA Kit. N=3 for each group. Neither 2μmol/L NY173 nor 30 μmol/L I942 have effect on the releasing of vWF from HUVECs. (C) Immunofluorescence of vWF in HUVECs. (a) Negative control, normal mouse IgGs were used as primary Abs. (b) NY173-treated, HUVECs were treated with 2μmol/L NY173 for 24 hrs. (c) I942-treated, HUVECs were treated with 30 μmol/L I942 for 24 hrs. In b and c, the samples were incubated with anti-vWF mouse monoclonal antibody as primary antibody, then stained with DAPI (blue) and Alexa Fluor 488- conjugated secondary antibody for vWF labels (green). Scale bars indicate 20 μm. The difference was not significant among the groups. (D) Quantitative analyses of vWF positive puncta and cell nuclei were performed using Image J software. The results were expressed as dot signals enumerated in each cell. Five microscopic fields were examined for each case. The results were expressed as dot signals enumerated in each cell. n = 3 for each group.

Figure S3: **Reverse transcription- quantitative polymerase chain reaction (RT-qPCR) analysis**

**of eNOS mRNA and iNOS mRNA expression in HUVECs.** (A)The effect of NY173 and I942 on eNOS mRNA expression in HUVECs. HUVECs were incubated with 2μmol/L NY173 or 30 μmol/L I942 for 24 hrs. N=3 for each group. It showed a non-significant trend towards lower eNOS mRNA expression in NY173 group and higher eNOS mRNA expression in I942 group comparing to vehicle. (B) The difference of iNOS mRNA expression among rTNFα group, NY173+rTNFα group and NY173+rTNFα+IRS group was not significant. n = 3 for each group.

Figure S4: **Model for EPAC1 regulating inflammation-triggered vWF releasing in the PI3K/eNOS-dependent manner.** Our proposed pathways from the present study are in black and previous reported pathways are in gray.

**Supplemental figure 1**

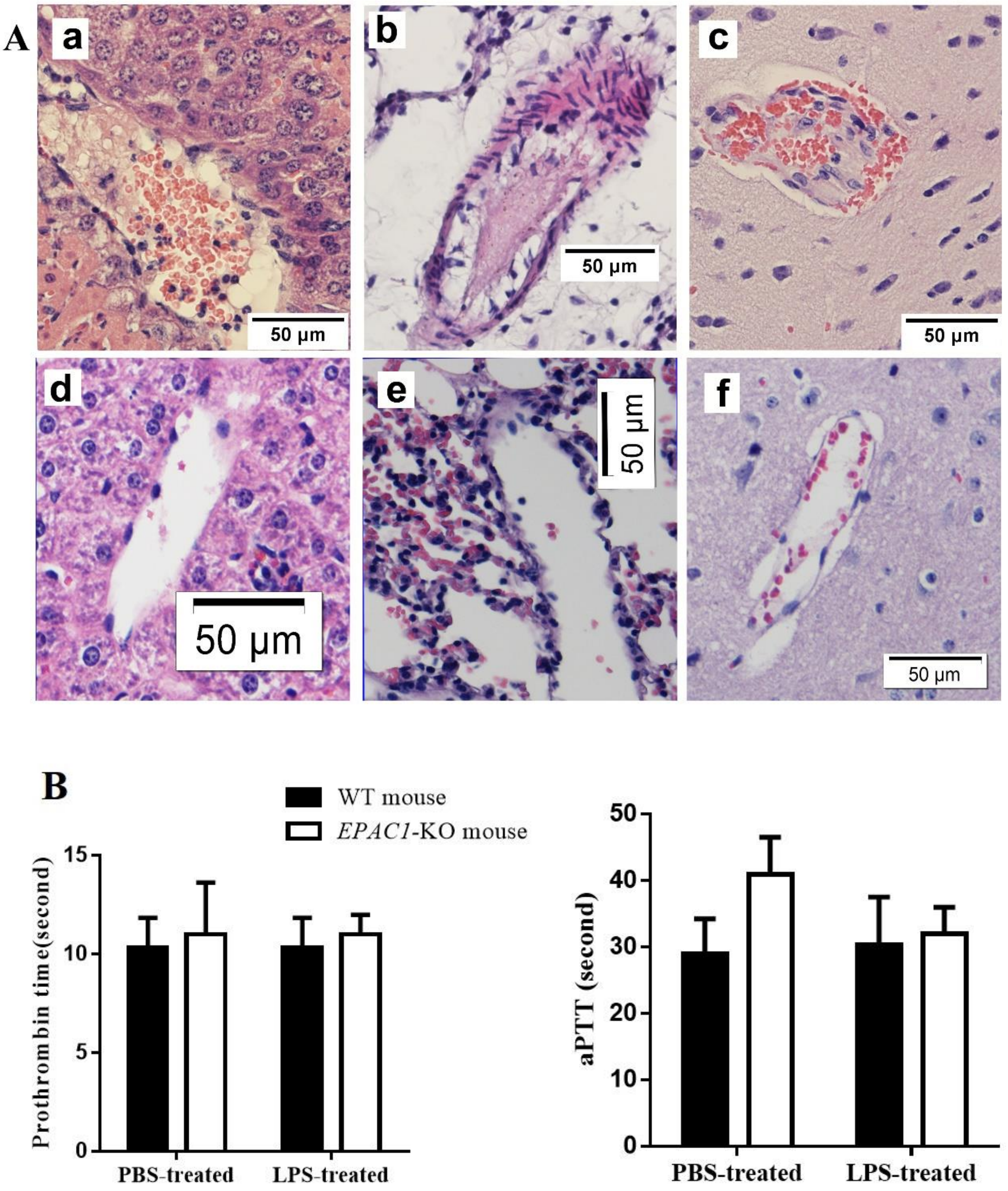

Supplemental figure 2

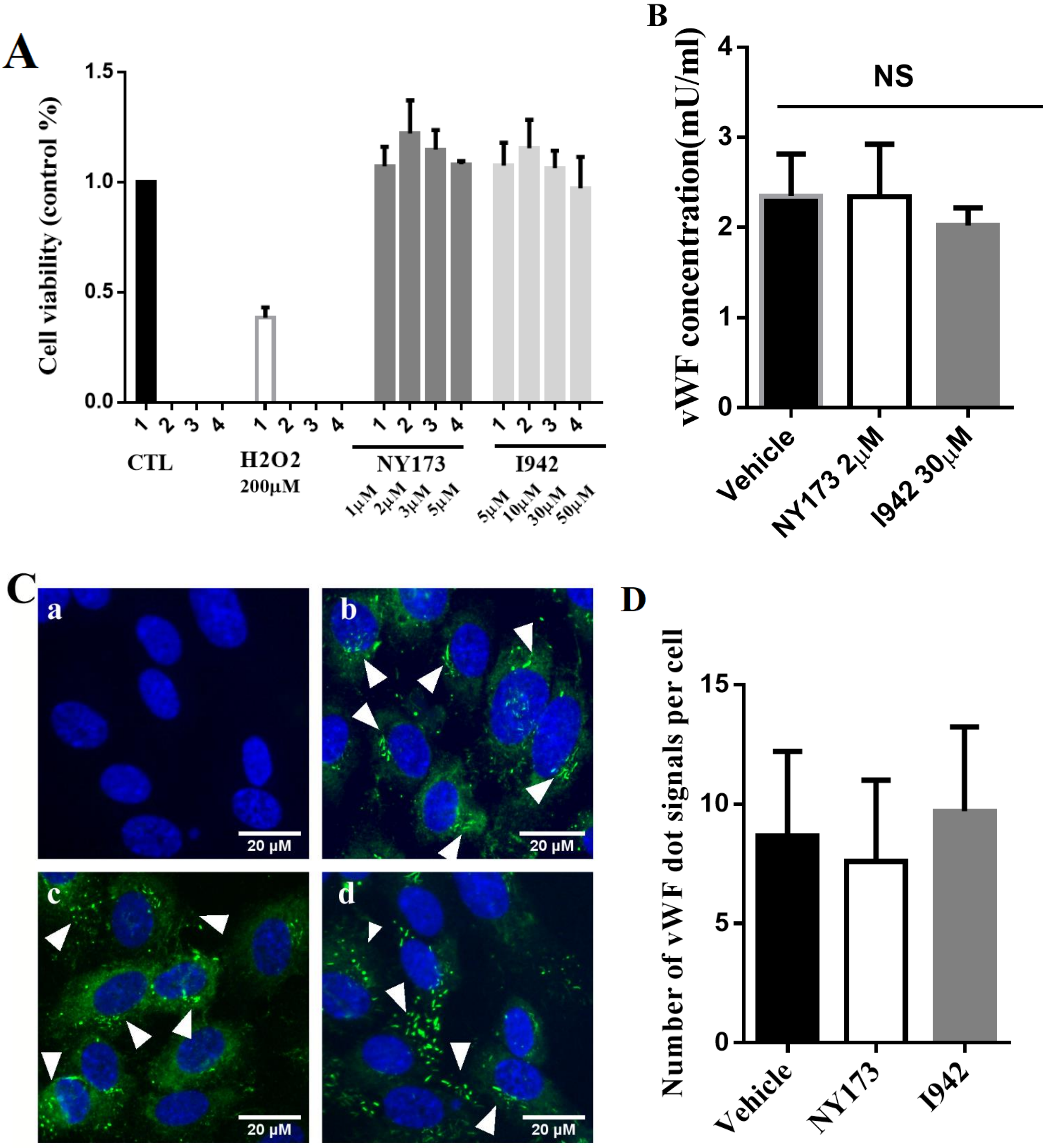

**Supplemental figure 3**

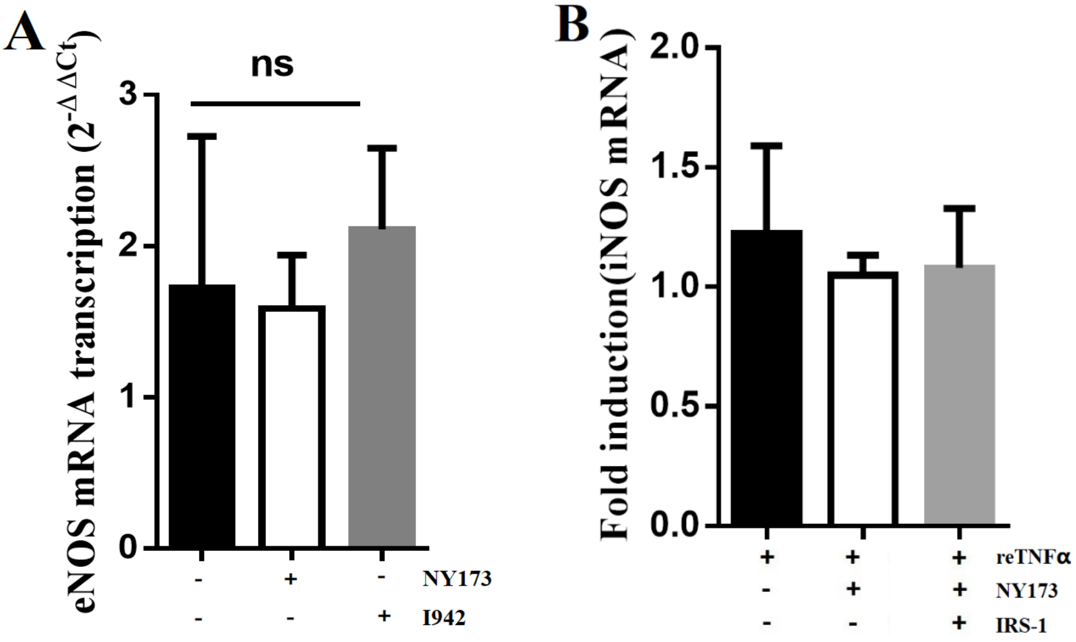

**Supplemental figure 4**

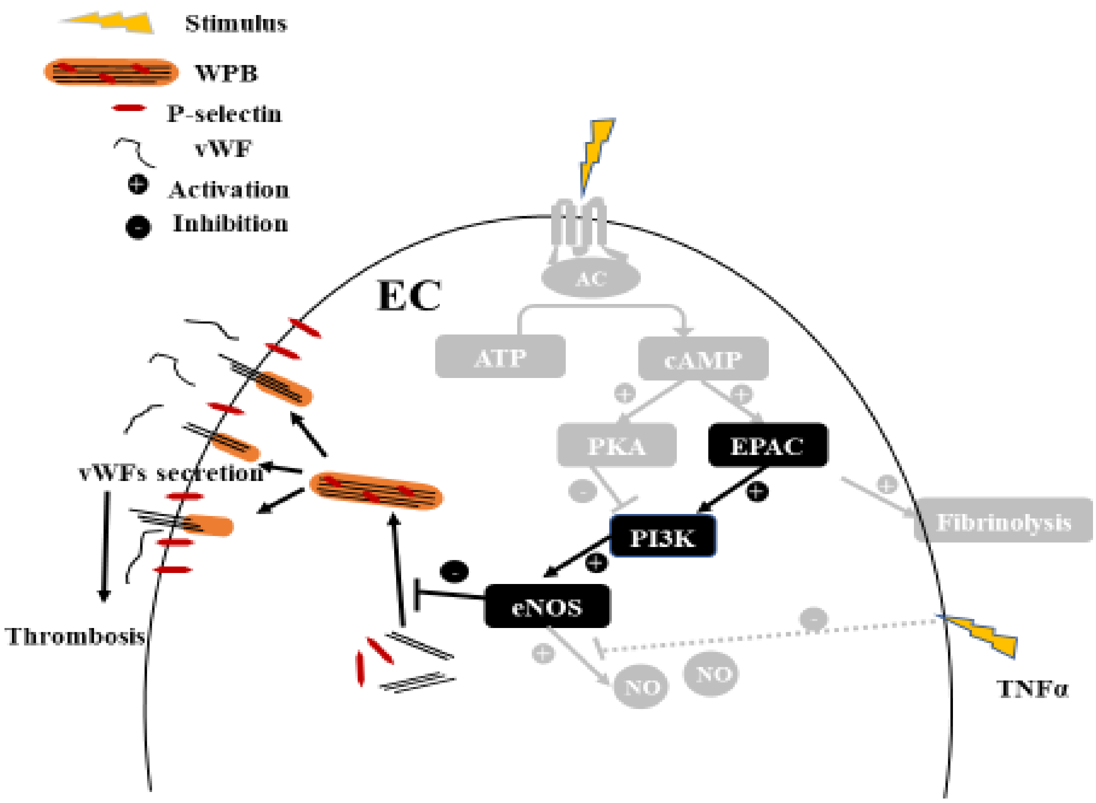
